# How does the host community structure affect the epidemiological dynamics of emerging infectious diseases?

**DOI:** 10.1101/2022.01.23.477158

**Authors:** Marina Voinson, Charline Smadi, Sylvain Billiard

## Abstract

Many pathogens, especially those responsible for emerging infectious diseases, are transmitted in a host community. How the host community structure affects an epidemic is still debated, particularly whether increasing the host community complexity would tend to amplify or dilute the incidence of an epidemic in a target population, *e*.*g*. humans or cattle. In this paper, we build a stochastic SIR model and compare epidemiological dynamics in a target population between three simple host community structures with an increasing complexity. Globally, our results show two possible main outcomes. First, an intermediate host can have a diluting effect by preventing the direct transmission from hosts to the target population, thus reducing the prevalence of infection. Second, when two sources of infection are considered, the effects of the epidemic are generally amplified. By highlighting that the structure of the ecological hosts network can dramatically affect epidemics, our results may have implications for the control of emerging infectious diseases.

## 1 Introduction

Zoonotic pathogens have a serious impact on socio-economic life in humans. More than 60% of human pathogens are classified as zoonoses [1, 2] i.e. pathogens coming from vertebrate animals. The pathogen either spreads efficiently in human populations once introduced from animals (e.g. influenza or human immunodeficiency virus), or spills over recurrently from an animal reservoir causing smaller outbreaks with high fatality rates (e.g. Nipah virus or Ebola virus). Emerging infectious diseases (EIDs) fall into the latter category of zoonoses. The pathogens responsible of the EIDs are naturally present in a reservoir in which they can live indefinitely [3] and emerge recurrently in human populations. 335 EIDs have been recorded since the 1940’s [4].

Many factors have been proposed to explain disease emergence including pathogen characteristics (e.g. mutation), characteristics of host populations (e.g. population size, migration) and ecological factors (e.g. land use, agriculture) [1, 5]. Human encroachment on wildlife habitats may also result in an increased transmission at the wildlife-human interface [6–8]. Moreover, the agriculture expansion and the increasing number of livestock can indirectly lead to an increasing emergence of diseases by acting as a bridge between humans and reservoir hosts. By changing their environment, human populations modify the contact rate between humans and potential reservoir hosts. Recent anthropogenic ecological change is likely to cause major changes to the geographic range leading to change in incidence of diseases by increasing the contact rate with the principal source of infection - animal populations [9].

Pathogens that can infect a broad range of hosts are ubiquitous and determining the source of infection is of primary interest to implement effective strategy of control [10]. The pathogen responsible of an emerging infectious disease and indefinitely maintained in an ecological system defines the reservoir [3, 11]. Human populations can be infected either by (i) a reservoir as the single source of infection; for instance, the emergence of the lassa virus is attributed to contacts with *Mastomys natalensis*, a common mouse in human households [12]. (ii) An intermediate host; the emerging infectious disease can infect a broad range of incidental hosts that are irrelevant for the long-term persistence of the pathogen. Then, the pathogen can spill over to the human population from those incidental populations and be the only source of infection. For instance, the infection by the hendra virus is due to a contact with a single intermediate host, the horses which are themselves infected by fruit bats, the reservoir [13]. (iii) Two sources of infection, i.e. a reservoir and one or several intermediate hosts. This is the case for the major part of pathogens responsible of EIDs such as Ebola virus, Marburg virus, Nipah and Middle East respiratory syndrom coronavirus (MERS) [14–17].

The effect of the cross-species transmission in the host community of a zoonotic disease remains poorly understood. Models tackling how cross-species spillovers affect zoonotic diseases are rare [18]. In a previous work, we showed that the size and frequency of outbreaks result from a complex interaction between the infectiousness of the disease and the spillover rate from a single reservoir [19]. However, most EIDs are involved in ecological networks with several sources of infection, *e*.*g*. one or several intermediate host species. Some authors have explored the effect of host-pathogen community assemblages (i.e. multi-host diversity) on epidemiological dynamics [20–22]. Two possible effects have been found, an amplifying effect in the case of density-dependent transmission and a buffering or dilution effect in the case of frequency-dependent transmission.

It is yet not clear whether the dilution vs. amplification effects are only due to the different transmission modes (frequency vs. density dependent) rather than on the variation of the host community structure. In addition, how dilution and amplification are defined and measured generally lacks accuracy and generality. Terms such as “dilution effect” (or “amplifying effect”) is used in a phenomenological sense to describe the situation when there is a decrease (or an increase) in disease frequency (i.e. the prevalence) [21], relatively to a reference model. In previous works, the effect of the multi-host diversity was explored with a single-host model as the reference, thus limiting the measurement of the amplification vs. dilution effects to a single event of infection [20–22]. Yet, EIDs are generally persistent in a community structure, which implies probable recurrent spillovers between host species. In such a situation, it is not known whether increasing the complexity of the host community structure tends to amplify or dilute the incidence of an epidemic in a target population.

In this paper, we propose to address the effect of the multi-host diversity by analyzing a stochastic model with a reservoir and two incidental populations. The pathogen is indefinitely maintained in the reservoir and can spill over to both incidental populations, the intermediate host and the target population. Our aim is to study the role of the intermediate host in addition to the reservoir on the epidemiological dynamics of the target population (e.g. the human population). In this multi-hosts process, five mechanisms can have an impact on the epidemiological dynamics of the target population, (1-2) the spillover transmission from the reservoir to both incidental population (i.e. the intermediate host and the target population), (3-4) the transmission between individuals within each incidental population and (5) the inter-incidental transmission, i.e. the transmission of the pathogen from the intermediate host to the target population. We addressed two main questions. What is the effect of the intermediate host on the epidemic disease observed in the target population and especially are intermediate hosts necessarily amplifiers of the infection as it is commonly thought? Second, what is the effect of the addition of a second source of infection? We show that depending on the host community structure considered, the intermediate host can have contrasted effects: it can dilute the epidemic disease when it is the only source of infection, while adding the intermediate host as a second source of infection in addition to the reservoir amplifies the epidemic disease.

## 2 Methods

### 2.1 The model

A model with different community structures is considered including a reservoir and two incidental populations, the intermediate host and the target population (Figure 1). The pathogen is assumed to live indefinitely in the reservoir and spills over recurrently into one or both incidental populations. We assume that the epidemic follows a stochastic SIR model (Susceptible Infected Recovered model) [23] in both incidental populations (see Figure 1). Each incidental population *x* is divided into three compartments: susceptible individuals (*S*_*x*_), infected individuals (*I*_*x*_) and recovered or dead individuals (*R*_*x*_). Individuals in the compartment *R*_*x*_ are not involved anymore in the transmission chains. The intermediate host and the population population will respectively be denoted with subscripts *h* and *p*.

**Figure 1:**
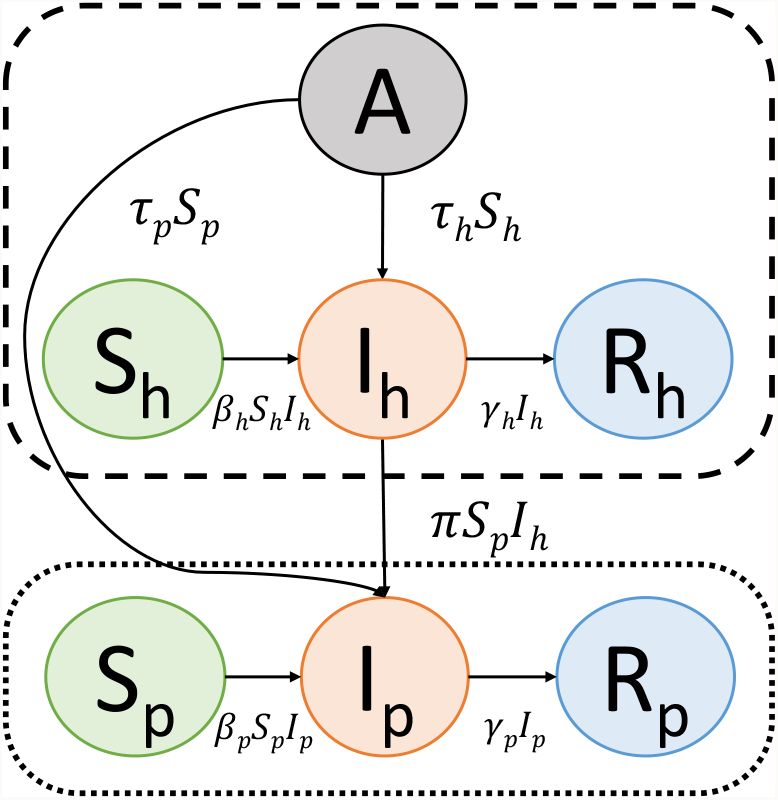
Interactions in the epidemiological community. The pathogen is persistent in the reservoir (*A*) and emerges recurrently in incidental populations at rate *τ*_*x*_*S*_*x*_. Two incidental populations are considered, the intermediate host and the target population (respectively subscripts *h* and *p*). In both incidental populations, the individuals are divided into three compartments depending on their epidemiological status (*S*_*x*_: susceptible, *I*_*x*_: infected and *R*_*x*_: recovered or dead individuals). A susceptible individual is infected by direct contact with an infected individual at rate *β*_*x*_*I*_*x*_ and an infected individual dies or recovers at rate *γ*_*x*_. A third way of infection is possible for the target population by a contact with an infected individual from the intermediate host (*I*_*h*_) at rate *πS*_*p*_*I*_*h*_. The target population is surrounded by a dotted line and the sources of infection for the target population are surrounded by a dashed line.

Two routes of transmission are considered: within- and between- populations. The pathogen is able to spread by inter-individual transmissions in both incidental populations as follows: a susceptible individual is infected by direct contact with an infected individual at rate *β*_*x*_ and recovers or dies at rate *γ*_*x*_. The basic reproductive ratio 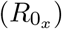 is equal to 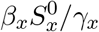 and represents the mean number of infected individuals that one infected individual generates during its infectious period in a whole susceptible population. 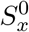 corresponds to the number of susceptible individuals at the beginning of the epidemic (at time *t* = 0). The incidental populations are connected to each other by the inter-incidental transmission which occurs at rate *π*. The pathogen from the intermediate host transmits to the target population by direct contact between an infected individual from the intermediate host (*I*_*h*_) and a susceptible individual from the target population (*S*_*p*_) at rate *πI*_*h*_*S*_*p*_. Moreover, the pathogen can emerge from the reservoir and spills over to the intermediate host at rate *τ*_*h*_ and over the target population at rate *τ*_*p*_.

The total number of susceptible individuals decreases during the epidemic since an individual cannot become susceptible after infection. We assume no reverse infections from the incidental populations to the reservoir and between incidental populations, which corresponds to many documented EID’s. For instance in the case of the Nipah virus, a fruit bat species is a reservoir host that maintains the pathogen indefinitely and transmits the pathogen to pigs [24]. While transmission may occur from bats to pigs and pigs to humans, there is no reciprocal transmissions from pigs to bats and from humans to pigs.

The demographic processes such as births and deaths are assumed to be much slower than the epidemiological ones, they are thus neglected. This is what is expected for an epidemic spreading locally during a short period of time. Three community structures will be compared. The first one is composed of the target population and the reservoir. The second and third ones are composed of the target population, the reservoir and the intermediate host species. (see Figure 2).

**Figure 2:**
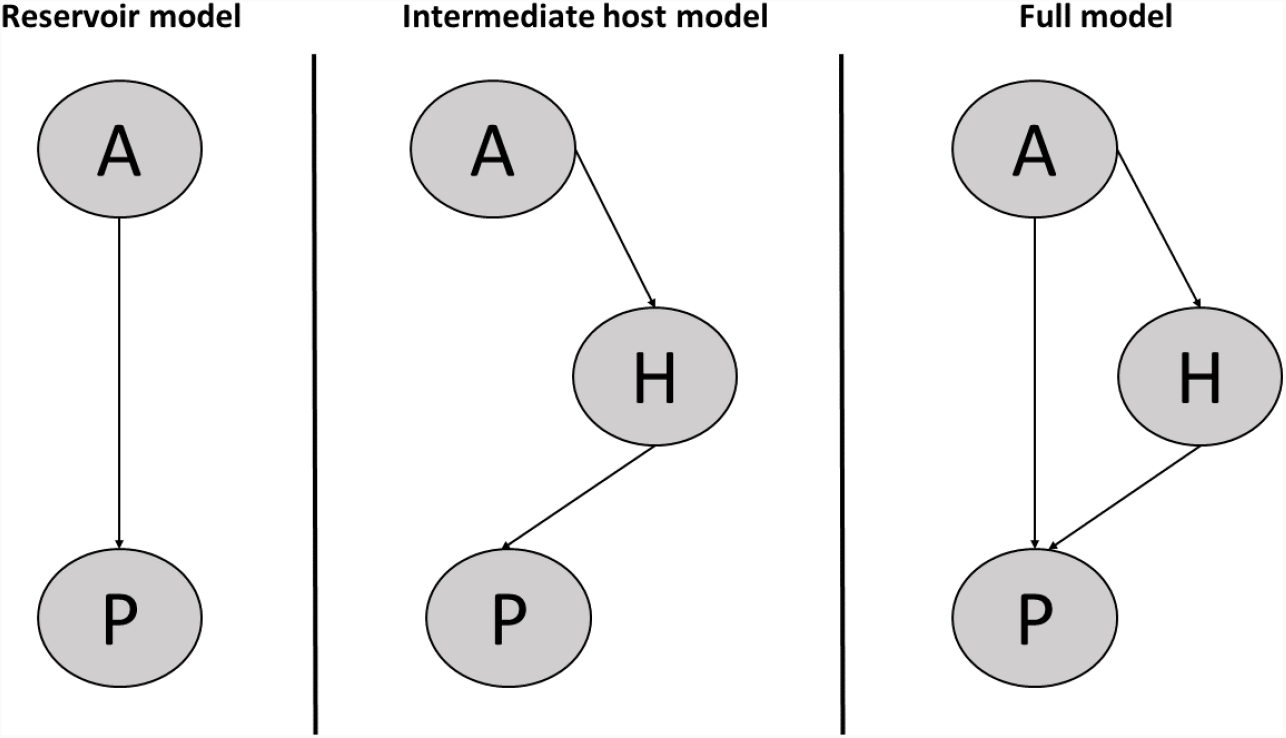
Community structure models. Left: most simple model, the only source of infection for the target population (P) is the reservoir (A), the pathogen spills over to the population at rate *τ*_*p*_*S*_*p*_. Middle: the only source of infection is the intermediate host (H) who is suffering from the infection by the reservoir (A) at rate *τ*_*h*_*S*_*h*_ and contaminates the target population (P) at rate *πS*_*p*_*I*_*h*_. Right: both the intermediate host (H) and the reservoir (A) infect the target population (P).

### 2.2 Simulation runs

The stochastic continuous time model presented in Figure 1 is analyzed with the help of numerical simulations obtained via a Gillespie algorithm described in Appendix A. The mathematical stochastic model with the reservoir as the single source of infection (Figure 2: the reservoir model), has been analyzed in a previous paper [19] where we showed that the size and frequency of outbreaks are affected by a complex interaction between the infectiousness of the disease and the spillover rate from the reservoir. The intermediate host and the target population are initially (*t* = 0) composed of 1000 susceptible individuals (*N*_*x*_ = *S*_*x*_ = 1000). Stochastic simulations were run for a basic reproductive ratio 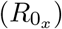 ranging from 0 to 4 and for a spillover (*τ*_*x*_) and an inter-incidental transmission rates (*π*) ranging from 10^−6^ to 10^−1^. The average number of outbreaks and the size of the largest outbreak have been measured in the target population. An outbreak occurs when the number of infected individuals reaches the epidemiological threshold (*I*_*p*_ ≥ *c*) and ends when there were no infected individuals anymore (*I*_*p*_ = 0). During one simulation multiple outbreaks can occur. In the case of non emerging infectious diseases, an epidemiological threshold is used to indicate the start of outbreaks. In the case of emerging infectious diseases, the value of the epidemiological threshold is very low because no infectious cases are expected. The value of *c* did not change qualitatively the results [19]. Simulations ended when there are no susceptible individuals anymore in the incidental populations.

### 2.3 Model analysis

In order to investigate the effect of the community structure, we compared the outcomes of an epidemic between three pairs of models differing by the complexity and the structure of the host community, thus highlighting different mechanisms (Figure 2 and table 2). The bridge effect: comparing the intermediate host model with the reservoir model where the intermediate host acts as a bridge between the reservoir and the target population, so the pathogen has to infect the intermediate host before spilling over to the target population. The intermediate host effect: an intermediate host is added as a second source of infection on the epidemic disease of the target population, whose effect will be measured by comparing the full model with the reservoir model. The reservoir effect: a reservoir is added as second source of infection, whose effect will be measured by comparing the full model with the intermediate host model.

**Table 1:** Parameters used and their values. UT denotes the unit of the time which can be expressed in days or weeks.

| Parameters | Values |
| --- | --- |
| $R_{0_x}$ | variable |
| $\gamma_x$ | $0.1 \text{ UT}^{-1}$ |
| $\tau_x$ | variable ($\text{UT}^{-1}$) |
| $\pi$ | variable ($\text{UT}^{-1}$) |
| $S_x$ | 1000 individuals |
| $c_x$ | 5 infected individuals |

**Table 2:**
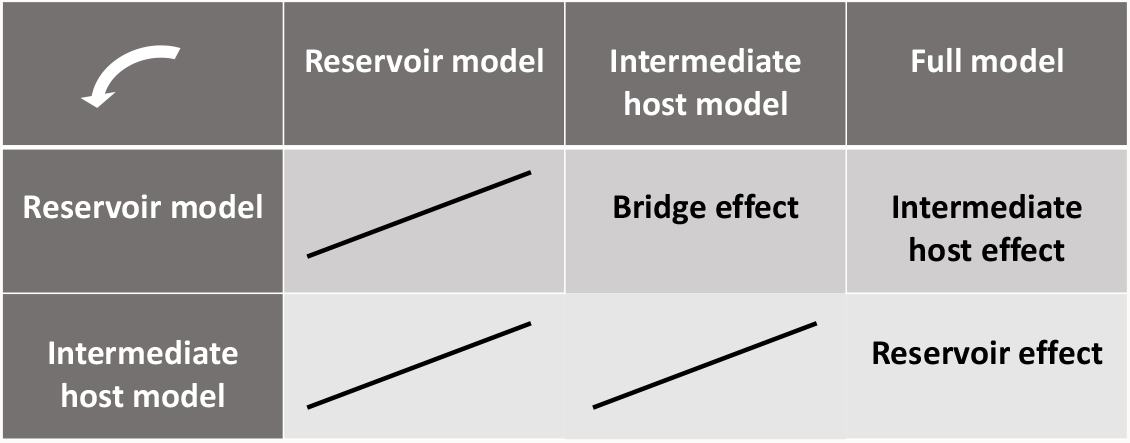
Mechanisms causing amplification or dilution as highlighted by model pairs comparison.

|  | Reservoir model | Intermediate host model | Full model |
| --- | --- | --- | --- |
| Reservoir model |  | Bridge effect | Intermediate host effect |
| Intermediate host model |  |  | Reservoir effect |

For each comparison (Table 2), three figures representing three qualitatively different dynamics for different parameters values are chosen. Results for other parameter sets are presented in Appendices B to D.

## 3 Results

### 3.1 Relative number of outbreaks

We measured how the number of outbreaks in the target population was affected by the host community structure when the basic reproductive number of the epidemic in the target population *R*0_*p*_ varies for different host community structures and inter-incidental transmission *π*. On Fig. 3, curves above the horizontal line at 1 show an amplification effect of the complexity of the community structure: the number of outbreaks is higher for the host community structure with the highest complexity and/or diversity. On the contrary, curves below the horizontal line at 1 show a dilution effect. Overall, Fig. 3 shows four different relationships between the host community structure and the basic reproductive number in the target population *R*0_*p*_. Increasing the complexity of the community can first show an amplification (e.g. Fig. 3i) or second a dilution (e.g. Fig. 3a) of the epidemic which decreases when the basic reproductive number 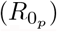 increases. Non-monotonous relationships are also observed with either a single maximum (third outcome, e.g. blue curves in Fig. 3g) or a minimum and a maximum (fourth outcome, e.g. blue curves in Fig. 3c). In the latter case, the minimum can be either above or below 1 which means that depending on the reproductive ration 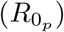 the host community structure can have either an amplification or a dilution effect (Fig. 3b and 3c).

**Figure 3:**
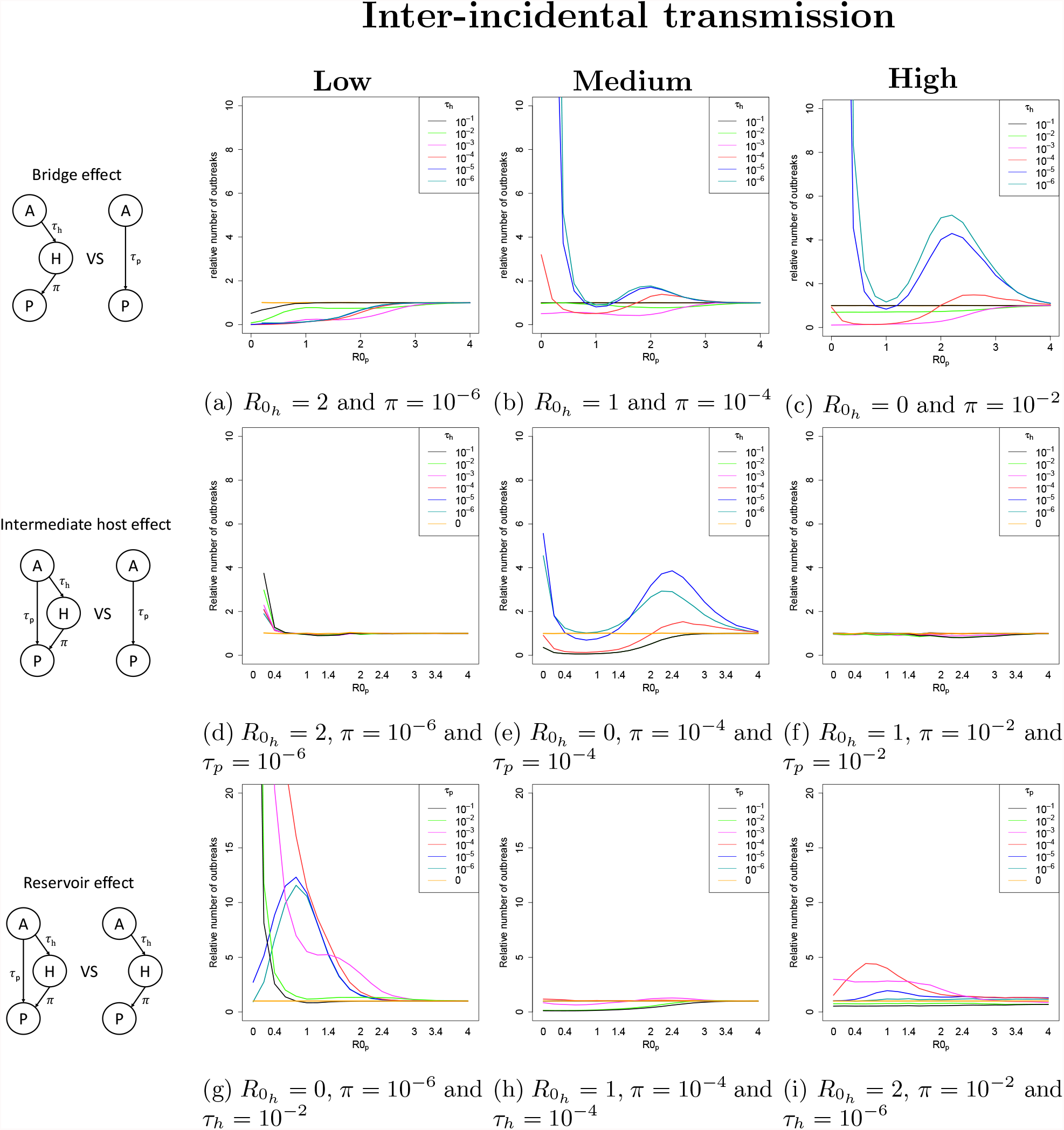
Relative number of outbreaks depending on the community structure. To analyze the bridge effect (subfigures a, b and c), the models are compared with *τ*_*p*_ = *τ*_*h*_.

Overall, our results show that there is a complex interaction between the host community structure and the basic reproductive number on an epidemic: only in some situations, increasing the complexity of the host community structure shows a unique effect. For instance, when the inter-incidental transmission is low, adding an intermediate host always shows dilution of the epidemic by decreasing the number of outbreaks (Fig. 3a), or always shows amplification when a transmission is added between the reservoir and the target population (Fig. 3g). In most situations, on the contrary, the community structure can amplify or dilute the epidemic depending on the inter-incidental transmission rate *π*, the basic reproductive number 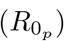 and the other transmission rates *τ*_*h*_ and *τ*_*p*_. Finally, the addition of an intermediate host can have a significant effect on the epidemic only when the inter-incidental transmission rate *π* is intermediate (Fig. 3e), and no significant when *π* is low or high (Fig. 3d and 3f).

Fewer outbreaks are observed when a second source of infection is added compared to the model with only one source of infection, either the intermediate host or the reservoir, and when the transmission rates within and between populations are high (Fig. 3e, 3f and Fig. 3h, 3i). Indeed, more frequent spillovers from the sources of infection (either the intermediate host or the reservoir or both), delay the extinction of the occurring outbreak in the target population and consume a higher proportion of the susceptible population. The next outbreak will then be less likely to reach the epidemiological threshold (*c*).

When the intermediate host acts as a bridge, adding a node can amplify epidemics when the inter-incidental transmission is intermediate or high and the spillover from the reservoir to the intermediate host is high because the consumption of the susceptible population in the intermediate host is low due to the fact that the infection will spread more slowly. Moreover, the recurrent emergence of the pathogen from the intermediate host to the target population will allow the outbreaks to reach the epidemiological threshold (*c*) more frequently in the target population when the inter-incidental transmission is intermediate or high (*π* ⩾ 10^−4^) (Figures 3b and 3c). Now if we consider the effect of the addition of a second source of infection (either the intermediate host or the reservoir), when the transmission rates are low, the second source of infection allows the small outbreaks in the target population to reach the epidemiological threshold *c* (Figures 3e, 3g and 3i).

### 3.2 Relative size of the largest outbreak

We also measured how the size of the largest outbreak in the target population was affected by the host community structure when the reproductive rate of the epidemic in the target population *R*0_*p*_ increases. Fig. 4 shows that the effect of the host community structure has simpler effect on the size of the largest outbreak than on the number of outbreaks (Fig. 3). Most often, the effect of the community structure has a monotonous relationship with the reproductive ratio 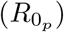.

**Figure 4:**
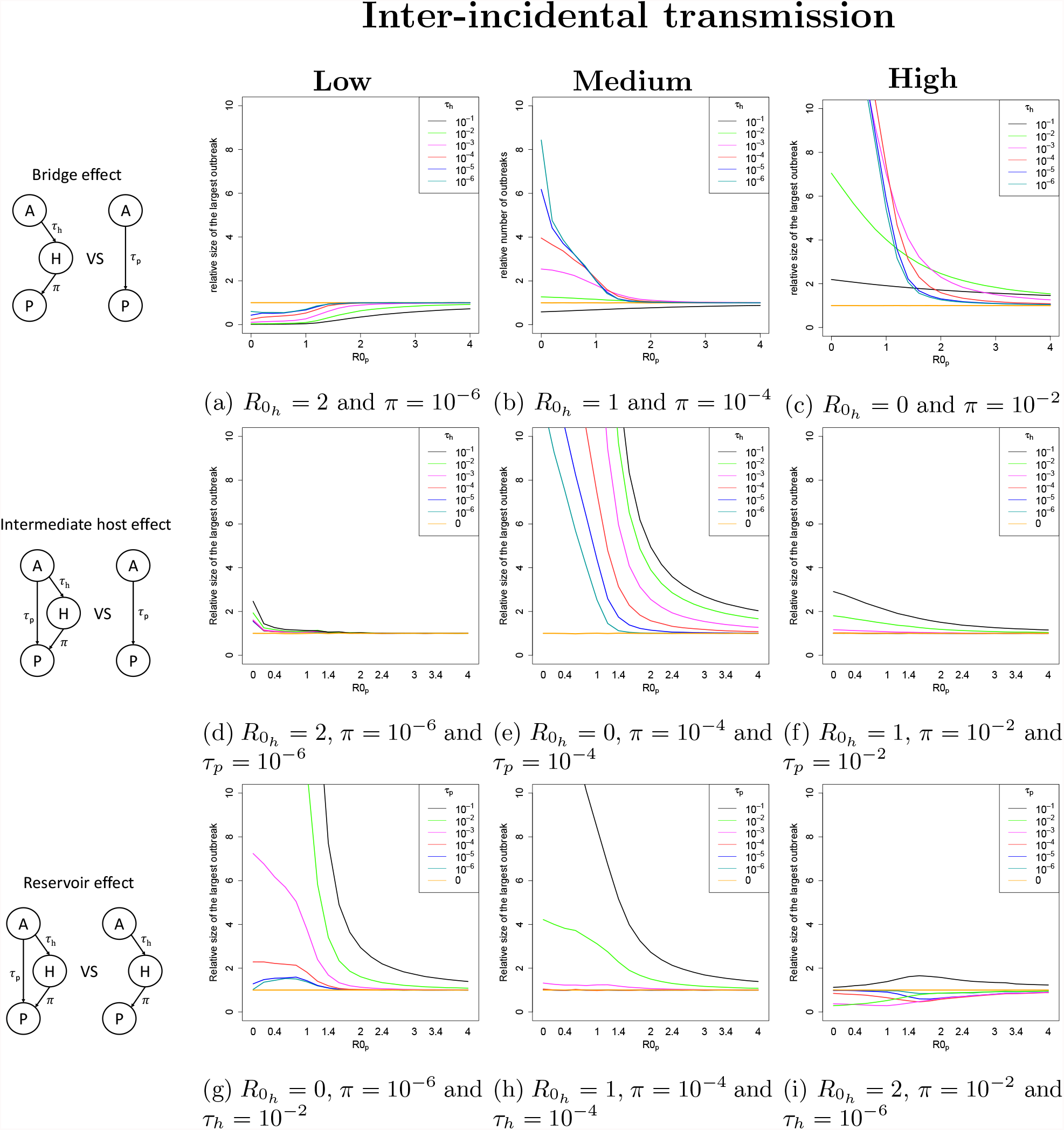
Relative size of the largest outbreak depending on the community structure. To analyze the bridge effect (subfigures a, b and c), the models are compared with *τ*_*p*_ = *τ*_*h*_.

Figure 4 shows that in general increasing the complexity of the community structure (i.e the addition of nodes and edges) acts as an amplifier for the size of outbreaks (with few exceptions). When the intermediate host acts as a bridge, the average size of the largest outbreak is always smaller when the inter-incidental transmission is low (*π* = 10^−6^) (see fig. 4a). Indeed, the infection propagates and consumes a large proportion of the susceptible population of the intermediate host (see Figure 5). Because the inter-incidental transmission depends on the number of infected individuals in the intermediate host, the size of the largest outbreak is smaller with the intermediate host than without. By contrast when the inter-incidental transmission is high (see Figures 4b and 4c), a higher average size of the largest outbreak is observed due to the high number of spillovers from the intermediate host to the target population. When a second source of infection is added, either the intermediate host or the reservoir, the largest outbreak becomes larger (see figs. 4d to 4i).

**Figure 5:**
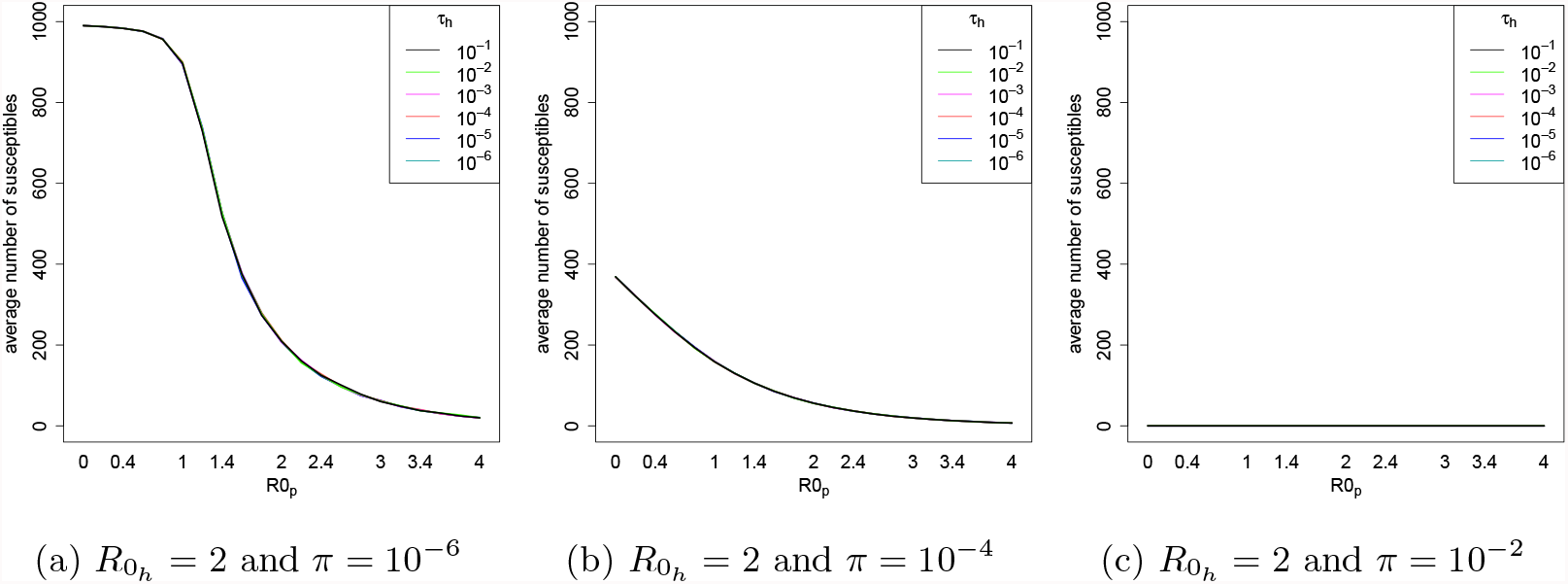
Average number of susceptible individuals remaining in the target population when there are only recovered individuals left in the intermediate host population. This average number is presented for the intermediate host model and is depicted as a function of the transmission rate between individuals in the target population 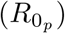. Different values of parameters have been used: a spillover rate (*τ*_*h*_) ranging from 10^−6^ to 10^−1^ and an inter-incidental transmission rate (*π*) equal to 10^−6^, 10^−4^ and 10^−2^.

### 3.3 Why does the intermediate host either amplify or dilute the disease incidence?

In order to address the effect of the intermediate host on the disease incidence, we collected data from our simulations about the remaining susceptible individuals in the target population (fig. 5). When the intermediate host is added between the reservoir and the target population, then a dilution of the epidemiological dynamics can be observed when the inter-incidental transmission is low. The outbreaks are then fewer and smaller. Indeed, the intermediate host is the only source of infection for the target population and the disease spreads between individuals by consuming the susceptibles in the intermediate population by contrast with an endemic reservoir. When the inter-incidental transmission is low, the intermediate host consumes only few susceptible individuals in the target population (see fig. 5). The ending of the disease epidemic in the intermediate host prevents the occurrence of outbreaks in the target population.

## 4 Discussion

Cross-species transmission is often associated with the emergence of pathogens in a broad range of hosts species and can have a serious impact on human life (e.g. the COVID-19 pandemic). Here, we studied how the community structure affects the number and the size of outbreaks in a target population (e.g. human population).

Using a simple SIR model with reservoir, our results showed that the addition of a second source of infection either amplifies or dilutes the epidemic disease depending on the statistics observed: the number of outbreaks tend to be lower while the size of the outbreaks is larger. In the case where human populations are already in contact with a reservoir, the addition of a second source of infection can amplify the severity of epidemics. Indeed, even when less outbreaks are expected, the outbreaks are larger and consume a larger proportion of susceptible individuals. This could explain observations during the 2013-2015 Ebola outbreak, the largest Ebola outbreak to date. Several factors have been proposed to explain this amplification of the epidemic disease such as a change in virulence, in the capacity of transmission between individuals or an increased contact between individuals [25, 26]. Here, we show that a larger size of outbreak could also be explained by a higher contact rate with the intermediate hosts, which are numerous in the case of Ebola virus (chimpanzees, gorillas, antelopes and bats) [27].

Broadly, a dilution effect of the number of outbreaks and the size of the largest outbreak is observed when the intermediate host acts as a bridge between the reservoir and the target population but only when the inter-incidental transmission is low. In this case, the number of outbreaks and the size of the largest outbreak are compared with the model with the reservoir as the only source of infection. This suggests that the within-host transmission in the intermediate host allows the dilution of the between-host transmission but only when this transmission is low. Other studies showed that in the case of a density-dependent transmission, the addition of species amplifies the disease in the target population [20, 21]. In those models, the epidemiological dynamics are compared with a single introduction in a SIR model and it is shown that the increasing host species diversity amplifies the frequency of the disease. In our case, the reference model used is the model with the reservoir acting as a source of infection. The pathogen spills over from the incidental population recurrently causing outbreaks and leading to a higher peak of infected individuals. The insertion of the intermediate host between the reservoir and the target host allows a lower number of outbreaks and a lower size for largest outbreak.

By analyzing the source of infection of a target population, a strategy of control can be implemented. In particular, when no drugs or other treatments are possible, zooprophylaxis can help decreasing human exposure to any given pathogen by using an animal population to divert the source of infection. This strategy of control has been implemented for malaria in order to divert insect blood feeding from humans to other animals, often cattle [28, 29]. The presence of multiple species can, in principle, have both a diluting effect, where the feeding on other species decreases the proportion of vectors feeding on the target species for a disease, and an amplifying effect where the access to multiple feeding hosts causes an increased abundance of vectors. When practicing zooprophylaxis, one assumes a dilution effect of animal vectors. In our model, a dilution effect is observed only when, instead of having a pathogen that spills over recurrently from an endemic reservoir, an intermediate host is placed between the endemic reservoir and the target population and acts as a bridge receiving the infection. When the contact between the intermediate host and human populations is low, and only in this case, the use of zooprophylaxis might be an interesting strategy of control since epidemiological dynamics in the intermediate host allows a decreasing of both the number of outbreaks and the size of the largest outbreak. Otherwise, if the reservoir infects the target population in addition to the intermediate host, even if weakly, then amplified epidemiological dynamics can be observed. Understanding the contact processes at the wildlife-human interface is mandatory before applying such an approach.

The outbreaks present in cattle or other animal populations in proximity with humans are empirically linked with an increasing number of infected individuals in humans. For instance outbreaks in pigs or in horses, caused respectively by the Nipah or Hendra viruses, are associated with the appearance of infected human cases [30]. Indeed, the Nipah and Hendra viruses propagate quickly and easily within non human animal conspecifics. Transmission between pigs in the same farm is attributed to direct contact with excretions and secretions such as urine, saliva, pharyngeal and lung secretions [24]. Once the pathogens are in pigs or horses, then they can emerge to humans especially to farmer workers. In our results we see that the increase of the within-population transmission increases slightly the peak of infected individuals in the target population but decreases the number of outbreaks. Indeed, a large number of individuals becomes infected in the intermediate hosts when the transmission within individuals is high leading to the rapid extinction of the outbreak. When an outbreak occurs in the intermediate host then an outbreak also emerges in the target population. The dynamics of the outbreak in the target population follows the dynamic of the outbreak in the intermediate host. In other words, when an outbreak starts in the intermediate hosts, an outbreak is also observed in the target population and on the contrary, when the outbreak stops in the intermediate hosts then the outbreak goes extinct in the target population. This phenomenon leads to a decreased number of outbreaks and a slightly increase of the size of the largest outbreak. We can conclude that the most important parameter that highly affects the epidemiological dynamics of the target population is the between-population transmission. Indeed, the contact rate between the target population and the intermediate host can increase both the peak of infected individuals and the number of outbreaks depending of the parameter values. The number of infected individuals is less important than the contact rate between-populations in the epidemiological dynamics. Those results can have an impact on strategy of control used. It seems more important to avoid contacts with the infected animal than to decrease the disease in the intermediate hosts.

To conclude, our results show that the addition of an intermediate host can have two opposite consequences. A dilution effect can be observed when the intermediate host is the only source of infection and prevents the direct contact between the reservoir and the target population, and an amplifying effect where the intermediate host acts as a second source of infection even when the pathogen emerges rarely from the reservoir to the target population.

**Appendices**

### A Algorithm of the full model

**Table A.1:** Transition rates of the stochastic model. The first three transitions correspond to the transitions of the classic SIR stochastic model for the intermediate host and the target population. The subscript *x* corresponds to subscripts *h* and *p* that identify respectively the intermediate host and the target population. Each epidemiological state is defined by *S*_*x*_: number of suceptible individuals; *I*_*x*_: number of infected individuals and *R*_*x*_: number of recovered individuals. *β*_*x*_ indicates the direct transmission rate. *τ*_*x*_ indicates the rate of spillover from the reservoir to both incidental populations and *π* corresponds to the inter-incidental transmission rate. Individuals recover at rate *γ*_*x*_.

| Transition |  | Effect | Transition rate |
| --- | --- | --- | --- |
| 1 | $(S_x, I_x, R_x) \rightarrow (S_x - 1, I_x + 1, R_x)$ | Direct transmission | $\beta_x S_x I_x$ |
| 2 | $(S_x, I_x, R_x) \rightarrow (S_x - 1, I_x + 1, R_x)$ | Spillover | $\tau_x S_x$ |
| 3 | $(S_x, I_x, R_x) \rightarrow (S_x, I_x - 1, R_x + 1)$ | Recovery | $\gamma_x I_x$ |
| 4 | $(S_p, I_p, R_p) \rightarrow (S_p - 1, I_p + 1, R_p)$ | Inter incidental transmission | $\pi S_p I_h$ |

The epidemiological dynamics of the target population using the full model (see Figure 2) can be simulated with the following algorithm (simulations were run in C++). At time *t*, knowing the population state, we can compute the total event rate, Ω (infections plus recoveries).

1. The total event rate Ω of the continuous time stochastic model is given by:

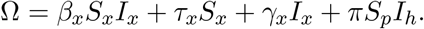
2. The next event time is *t*′ = *t* + *δ* where *δ* is exponentially distributed with parameter Ω.
3. The next event to occur is randomly chosen: direct transmission, spillover transmission, recover or inter incidental transmission with respective probabilities *β*_*x*_*S*_*x*_*I*_*x*_/Ω, *τ*_*x*_*S*_*x*_*/*Ω, *γ*_*x*_*I*_*x*_/Ω and *πS*_*p*_*I*_*h*_/Ω.

We performed stochastic individual-based simulations of the epidemics with spillover transmission, using rates as presented in Table A.1. The incidental host is initially (*t* = 0) composed of 1000 susceptible individuals (*N*_*x*_ = *S*_*x*_ = 1000). The infection is considered as endemic in the reservoir. Simulations are stopped when there is no susceptible individual anymore. An outbreak begins when the number of infected individuals reaches the epidemiological threshold *c* (*c* = 5 infected individuals in the simulations) and ends when in the target population there is no infected individual anymore (*I*_*p*_ = 0). Stochastic simulations were run for values of the basic reproductive ratio (*R*_0_) ranging from 0 to 4 and of the spillover transmission (*τ*_*x*_) and the inter-incidental transmission (*π*) ranging from 10^−6^ to 10^−1^. 1000 simulations are performed for each parameter set. All other parameter values are detailed in Table 1.

### B. The intermediate host model compared to the reservoir model

#### B.1 The relative number of outbreaks

**Figure B.1:**
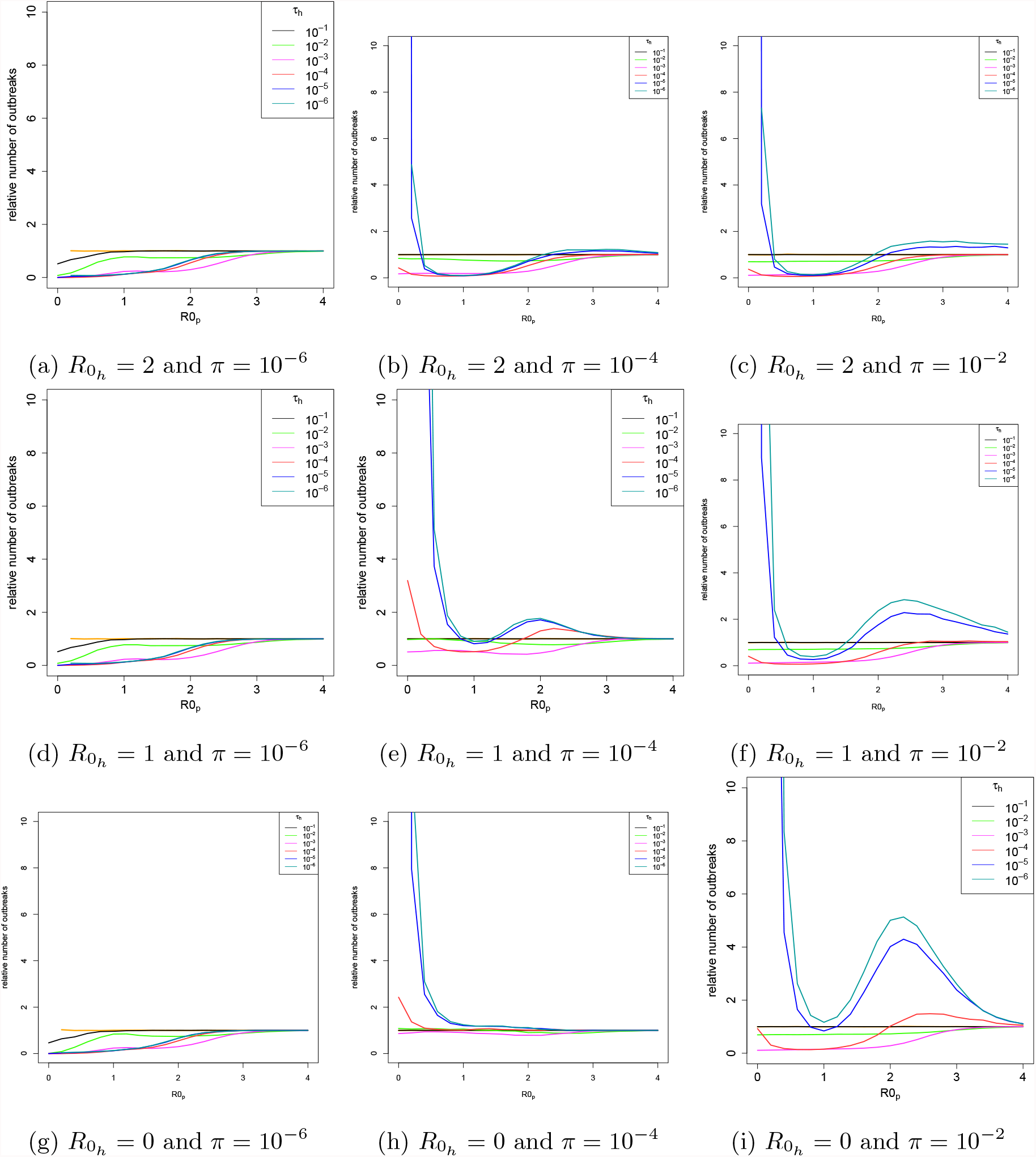
Relative number of outbreaks in the target population in the intermediate host model compared to the reservoir model. A low, intermediate and high value of the within-host transmission in the intermediate host (respectively 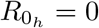, 1 and 2) and inter-incidental transmission (respectively *π* = 10^−6^, 10^−4^ and 10^−2^) is considered. The relative number of outbreaks is depicted as a function of the direct transmission between individuals 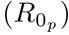. Each curve represents a spillover transmission from the reservoir to the intermediate host (*τ*_*h*_) ranging from 10^−6^ to 10^−1^.

#### B.2 The relative size of the largest outbreak

**Figure B.2:**
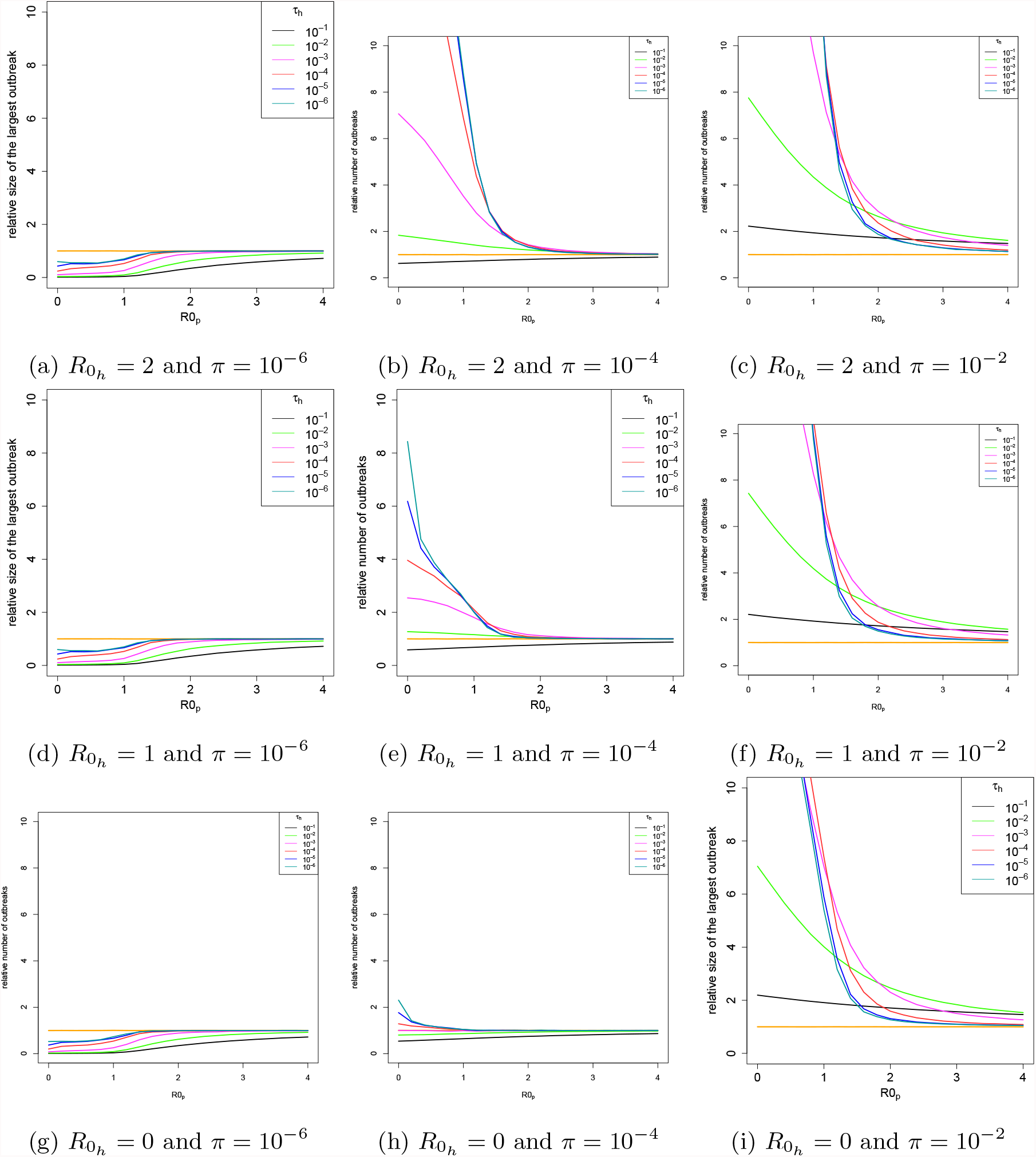
Relative size of the largest outbreak in the target population in the intermediate host model compared to the reservoir model. A low, intermediate and high value of the within-host transmission in the intermediate host (respectively 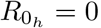, 1 and 2) and inter-incidental transmission (respectively *π* = 10^−6^, 10^−4^ and 10^−2^) is considered. The relative size of the largest outbreak is depicted as a function of the direct transmission between individuals 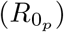. Each curve represents a spillover transmission from the reservoir to the intermediate host (*τ*_*h*_) ranging from 10^−6^ to 10^−1^.

### C. The full model compared to the reservoir model

#### C.1 The relative number of outbreaks

**Figure C.1:**
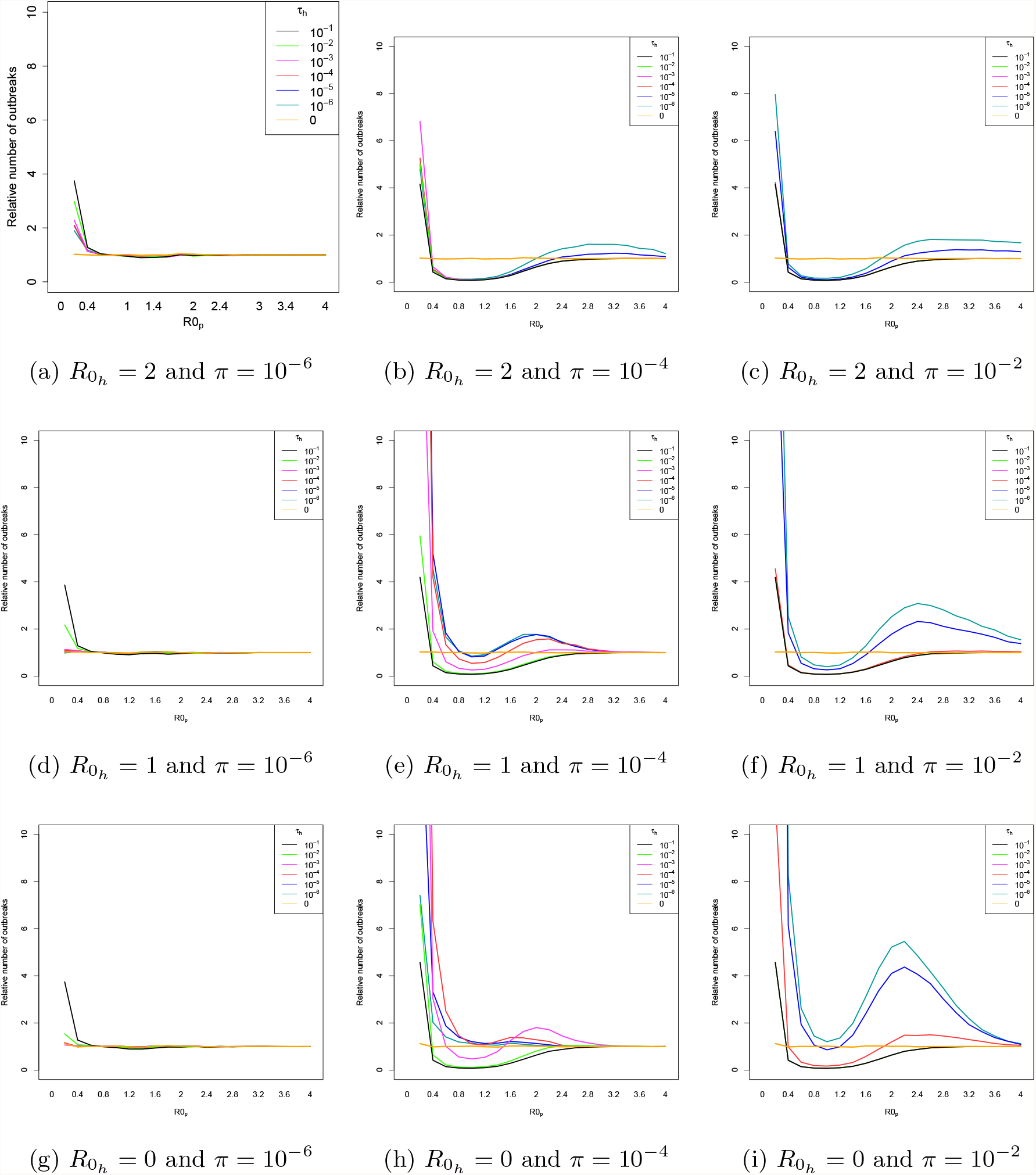
Relative number of outbreaks between the full model and the reservoir model. A low, intermediate and high value of the transmission between individuals in the intermediate host (respectively 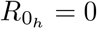, 1 and 2) and inter-incidental transmission (respectively *π* = 10^−6^, 10^−4^ and 10^−2^) are considered. The relative number of outbreaks is depicted as a function of the transmission between individuals in the target population 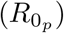. Each curve represents a spillover transmission from the reservoir to the intermediate host (*τ*_*h*_) ranging from 10^−6^ to 10^−1^. A low effect of the reservoir is considered (*τ*_*p*_ = 10^−6^).

**Figure C.2:**
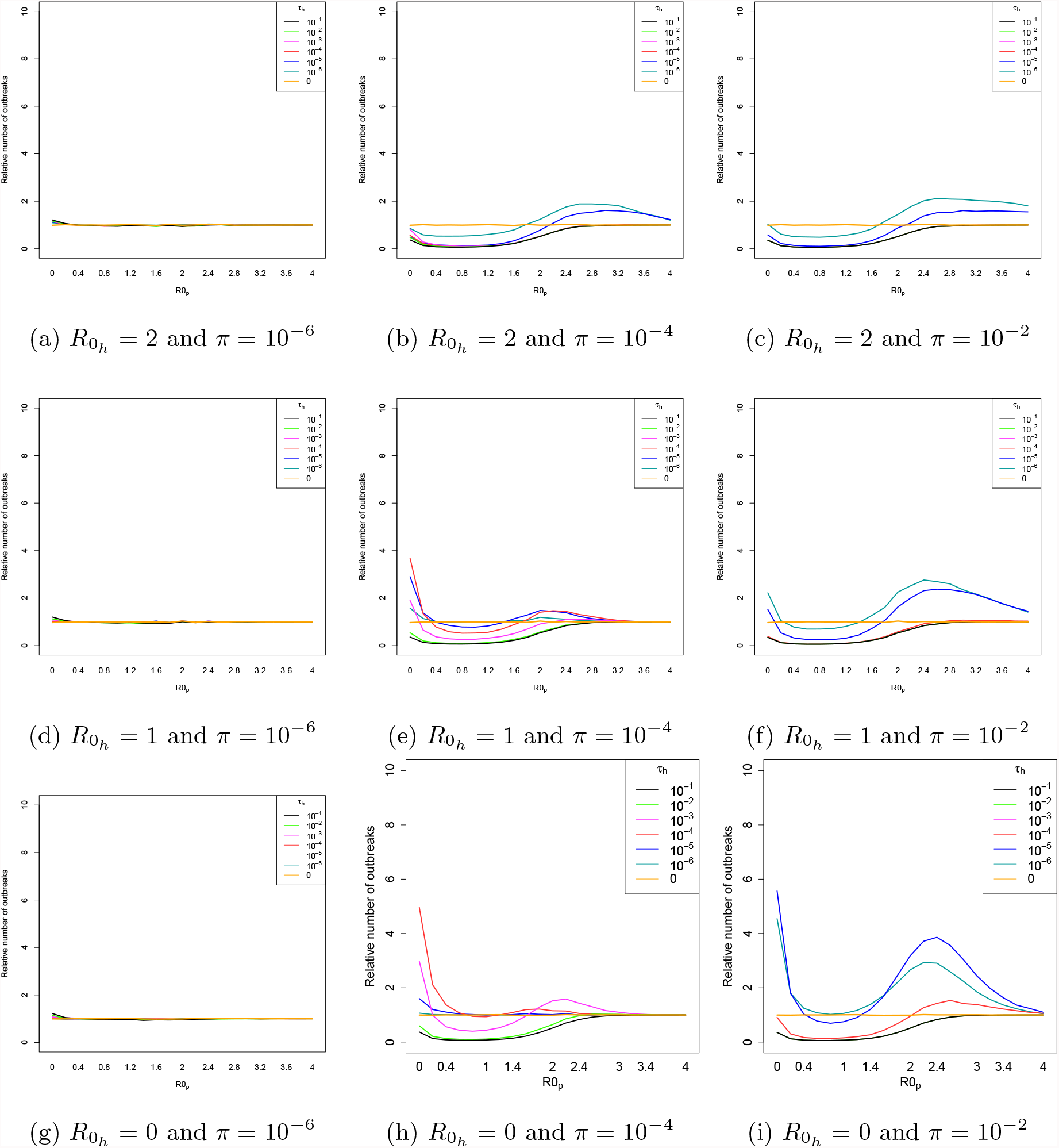
Relative number of outbreaks between the full model and the reservoir model. A low, intermediate and high value of the transmission between individuals in the intermediate host (respectively 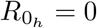, 1 and 2) and inter-incidental transmission (respectively *π* = 10^−6^, 10^−4^ and 10^−2^) are considered. The relative number of outbreaks is depicted as a function of the transmission between individuals in the target population 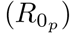. Each curve represents a spillover transmission from the reservoir to the intermediate host (*τ*_*h*_) ranging from 10^−6^ to 10^−1^. An intermediate effect of the reservoir is considered (*τ*_*p*_ = 10^−4^).

**Figure C.3:**
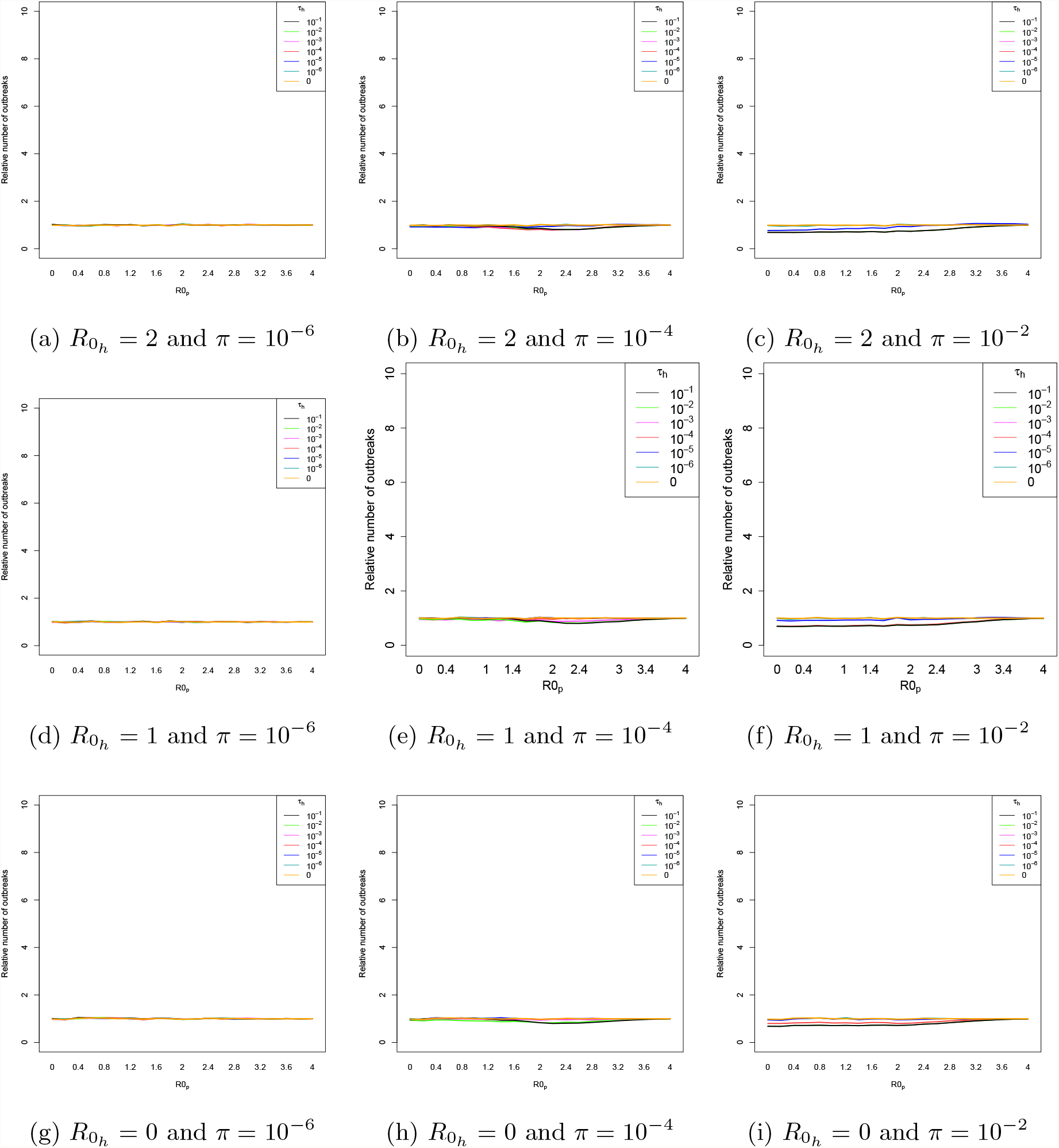
Relative number of outbreaks between the full model and the reservoir model. A low, intermediate and high value of the transmission between individuals in the intermediate host (respectively 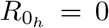, 1 and 2) and inter-incidental transmission (respectively *π* = 10^−6^, 10^−4^ and 10^−2^) are considered. The relative number of outbreaks is depicted as a function of the transmission between individuals in the target population 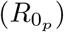. Each curve represents a spillover transmission from the reservoir to the intermediate host (*τ*_*h*_) ranging from 10^−6^ to 10^−1^. A high effect of the reservoir is considered (*τ*_*p*_ = 10^−2^).

#### C.2 The relative size of the largest outbreak

**Figure C.4:**
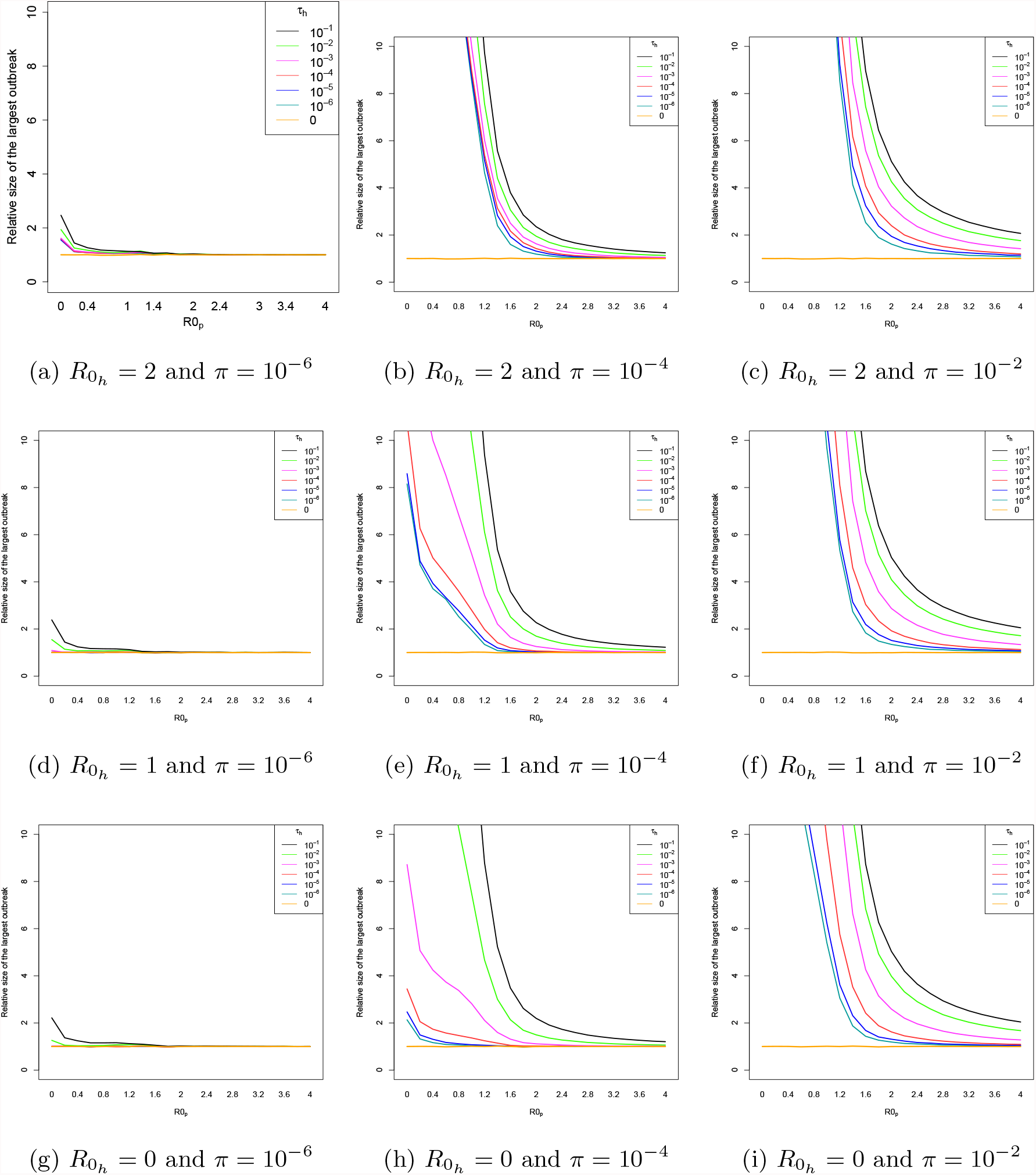
Relative size of the largest outbreak between the full model and the reservoir model. A low, intermediate and high value of the transmission between individuals in the intermediate host (respectively 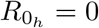, 1 and 2) and inter-incidental transmission (respectively *π* = 10^−6^, 10^−4^ and 10^−2^) are considered. The relative size of the largest outbreak is depicted as a function of the transmission between individuals in the target population 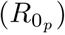. Each curve represents a spillover transmission from the reservoir to the intermediate host (*τ*_*h*_) ranging from 10^−6^ to 10^−1^. A low effect of the reservoir is considered (*τ*_*p*_ = 10^−6^).

**Figure C.5:**
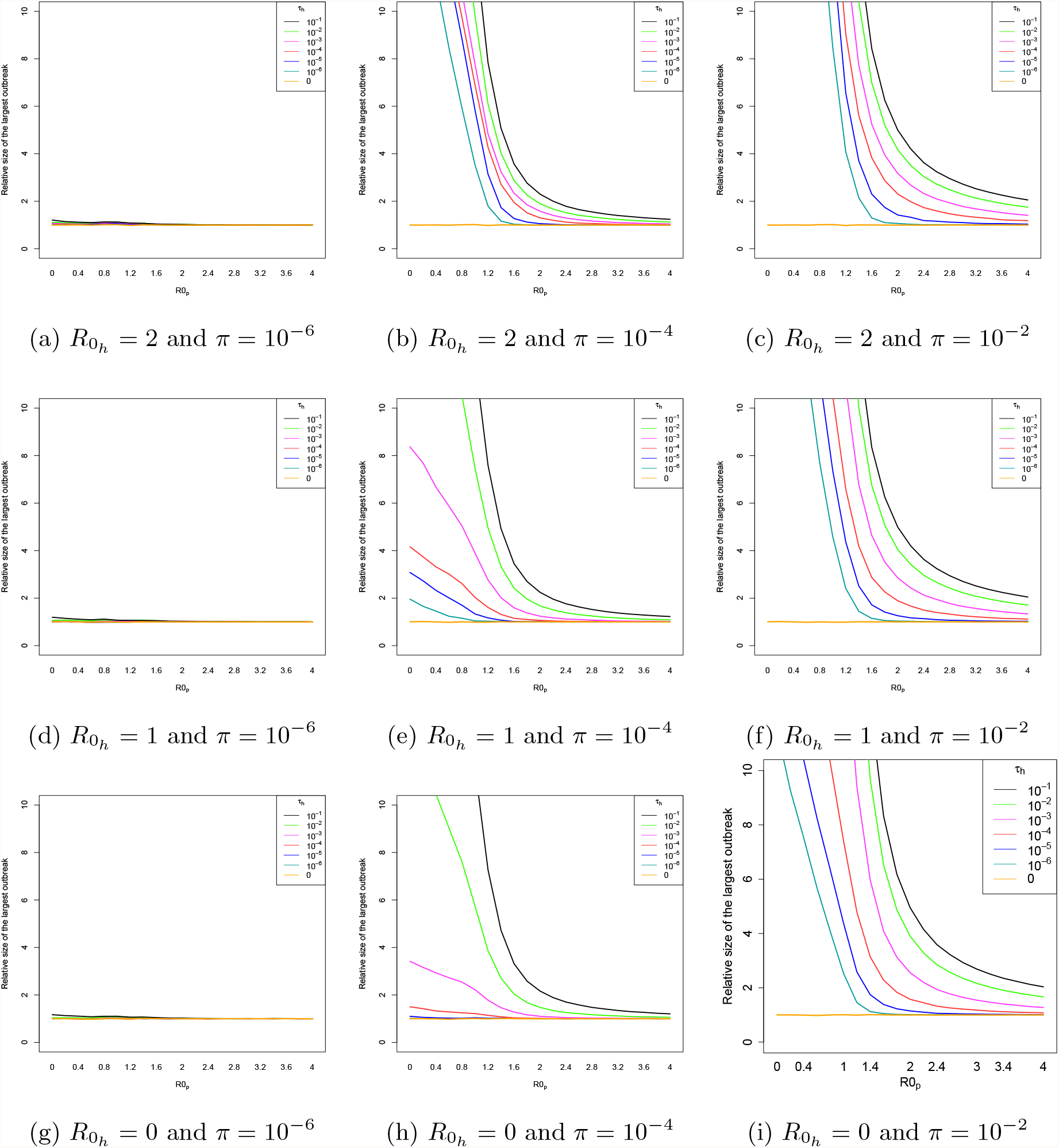
Relative size of the largest outbreak between the full model and the reservoir model. A low, intermediate and high value of the transmission between individuals in the intermediate host (respectively 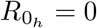, 1 and 2) and inter-incidental transmission (respectively *π* = 10^−6^, 10^−4^ and 10^−2^) are considered. The relative size of the largest outbreak is depicted as a function of the transmission between individuals in the target population 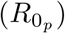. Each curve represents a spillover transmission from the reservoir to the intermediate host (*τ*_*h*_) ranging from 10^−6^ to 10^−1^. An intermediate effect of the reservoir is considered (*τ*_*p*_ = 10^−4^).

**Figure C.6.**
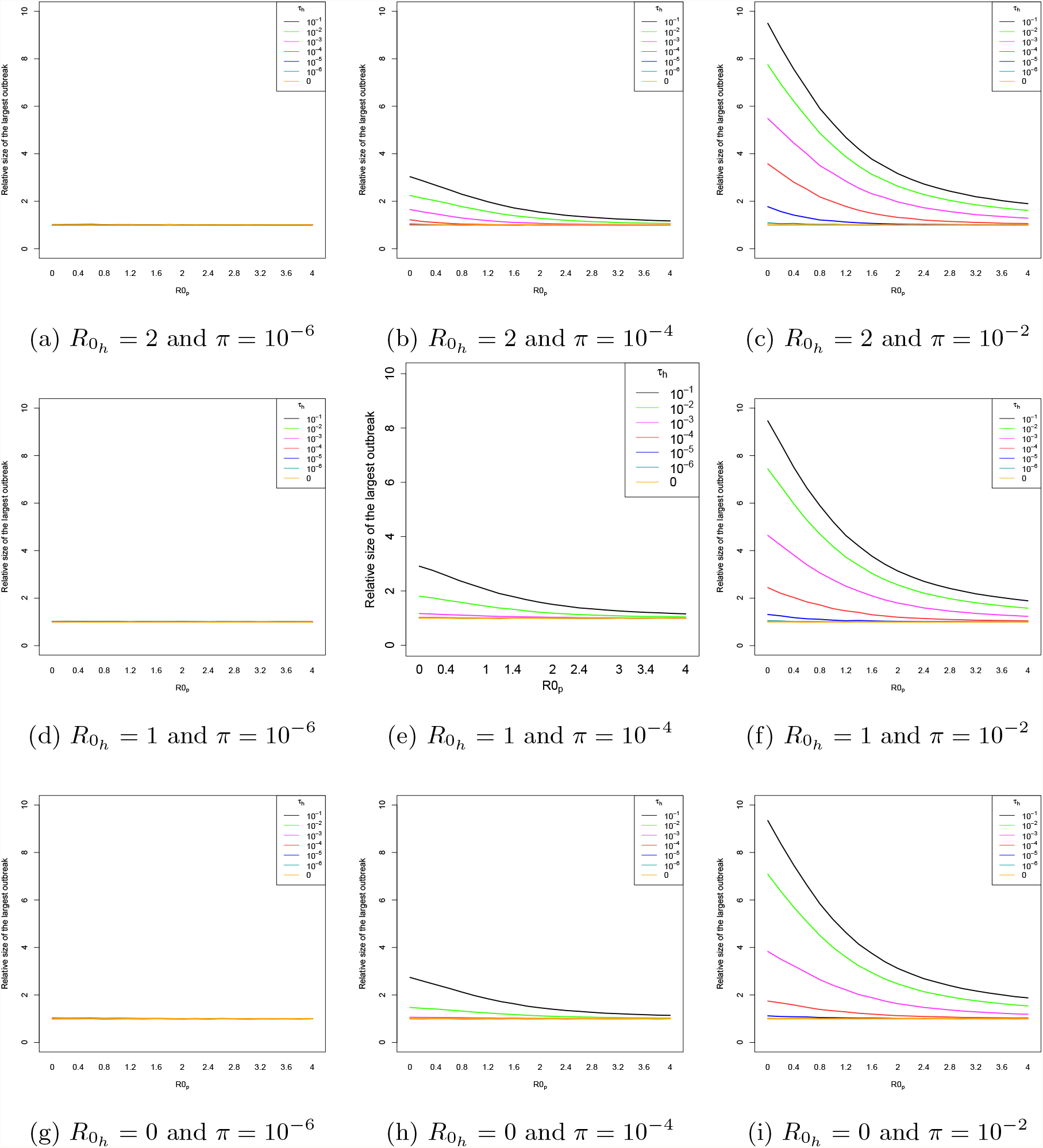
Relative size of the largest outbreak between the full model and the reservoir model. A low, intermediate and high value of the transmission between individuals in the intermediate host (respectively 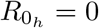, 1 and 2) and inter-incidental transmission (respectively *π* = 10^−6^, 10^−4^ and 10^−2^) are considered. The relative size of the largest outbreak is depicted as a function of the transmission between individuals in the target population 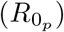. Each curve represents a spillover transmission from the reservoir to the intermediate host (*τ*_*h*_) ranging from 10^−6^ to 10^−1^. A high effect of the reservoir is considered (*τ*_*p*_ = 10^−2^).

### D The full model compared to the intermediate host model

#### D.1 The relative number of outbreaks

**Figure D.1:**
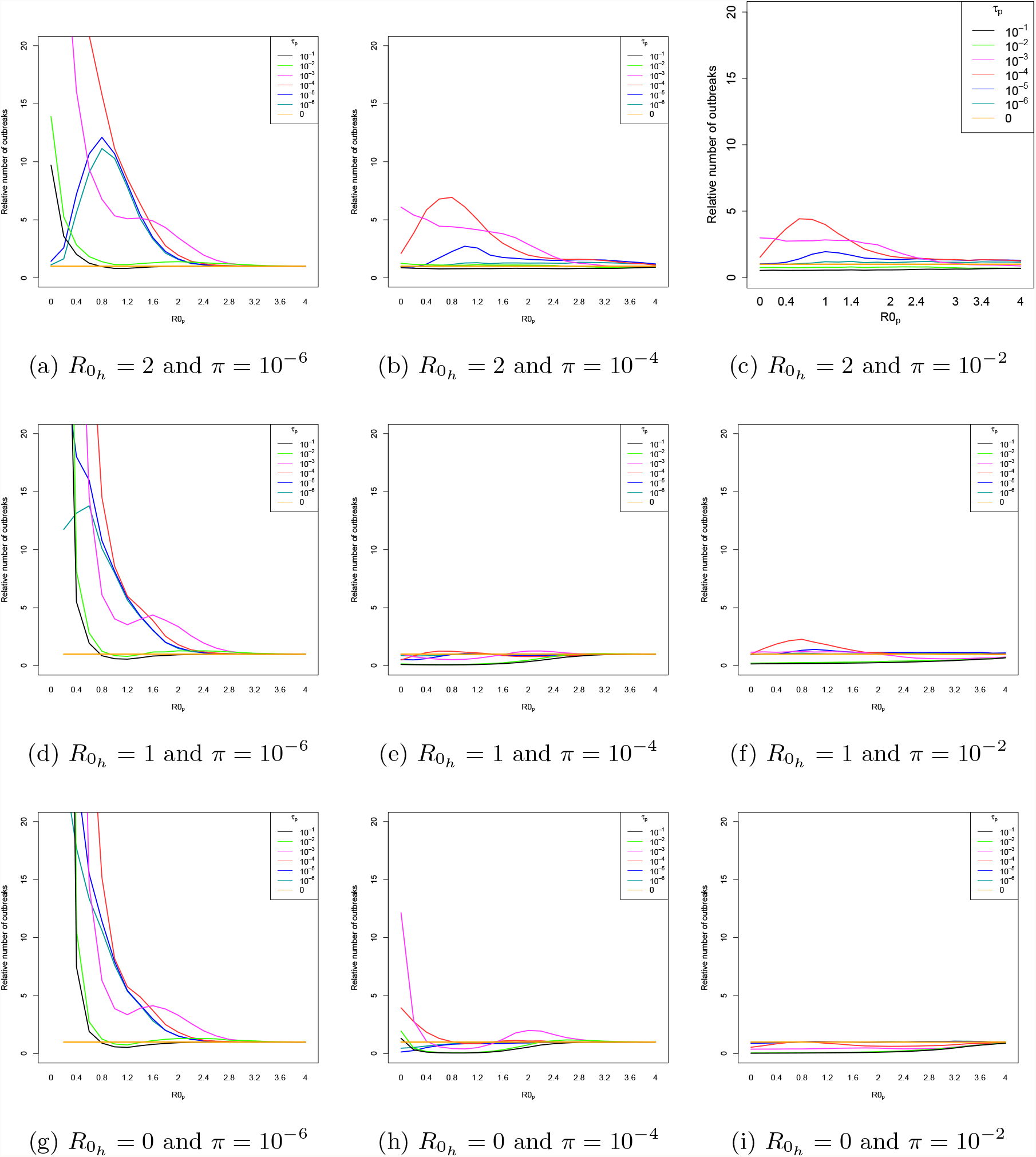
Relative number of outbreaks between the full model and the reservoir model. A low, intermediate and high value of the transmission between individuals in the intermediate host (respectively 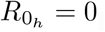, 1 and 2) and inter-incidental transmission (respectively *π* = 10^−6^, 10^−4^ and 10^−2^) are considered. The relative number of outbreaks is depicted as a function of the transmission between individuals in the target population 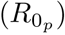. Each curve represents a spillover transmission from the reservoir to the target population (*τ*_*p*_) ranging from 10^−6^ to 10^−1^. A low spillover rate from the reservoir to the intermediate host is considered *τ*_*h*_ = 10^−6^.

**Figure D.2:**
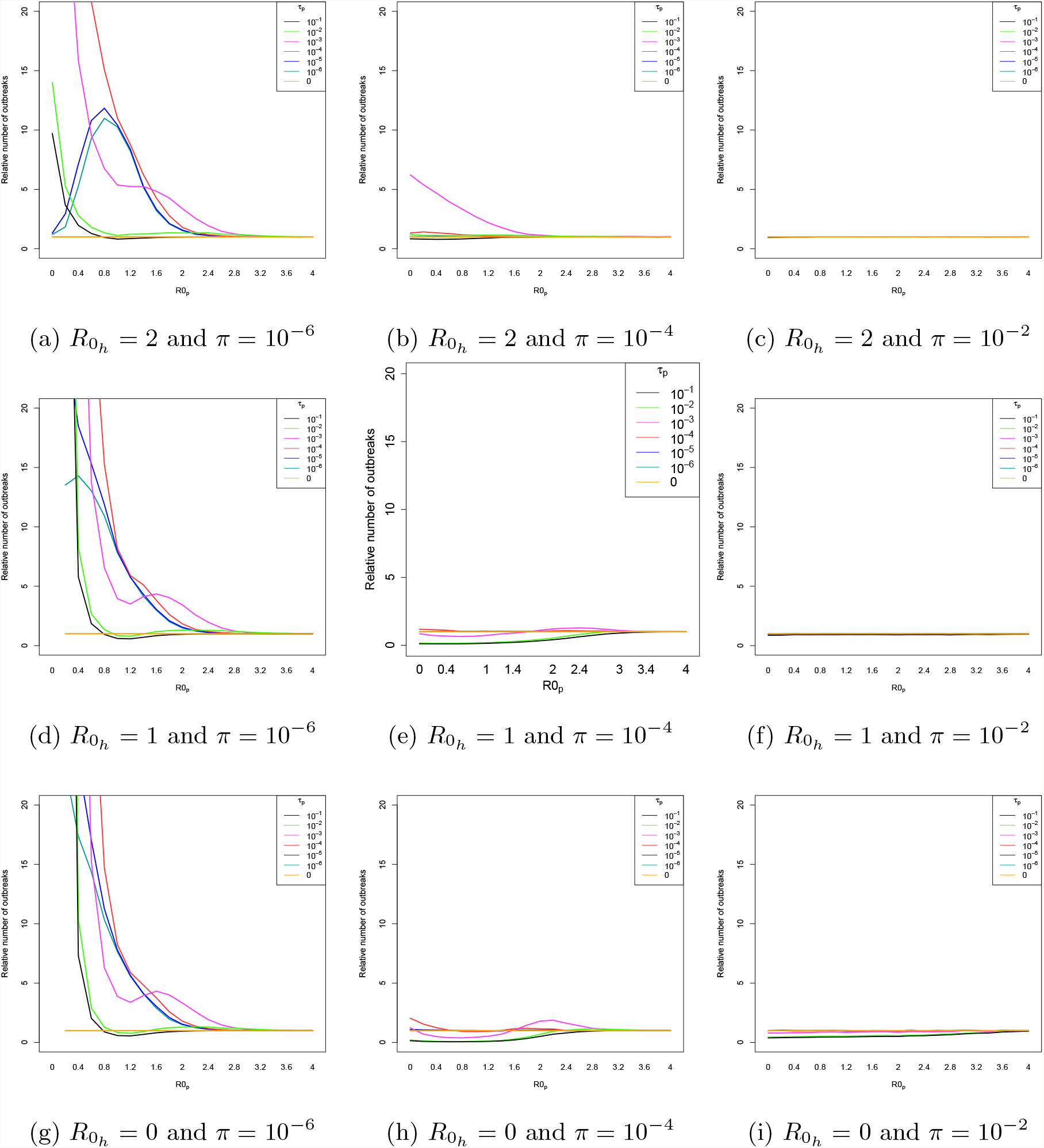
Relative number of outbreaks between the full model and the reservoir model. A low, intermediate and high value of the transmission between individuals in the intermediate host (respectively 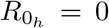, 1 and 2) and inter-incidental transmission (respectively *π* = 10^−6^, 10^−4^ and 10^−2^) are considered. The relative number of outbreaks is depicted as a function of the transmission between individuals in the target population 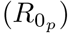. Each curve represents a spillover transmission from the reservoir to the target population (*τ*_*p*_) ranging from 10^−6^ to 10^−1^. An intermediate spillover rate from the reservoir to the intermediate host is considered *τ*_*h*_ = 10^−4^.

**Figure D.3:**
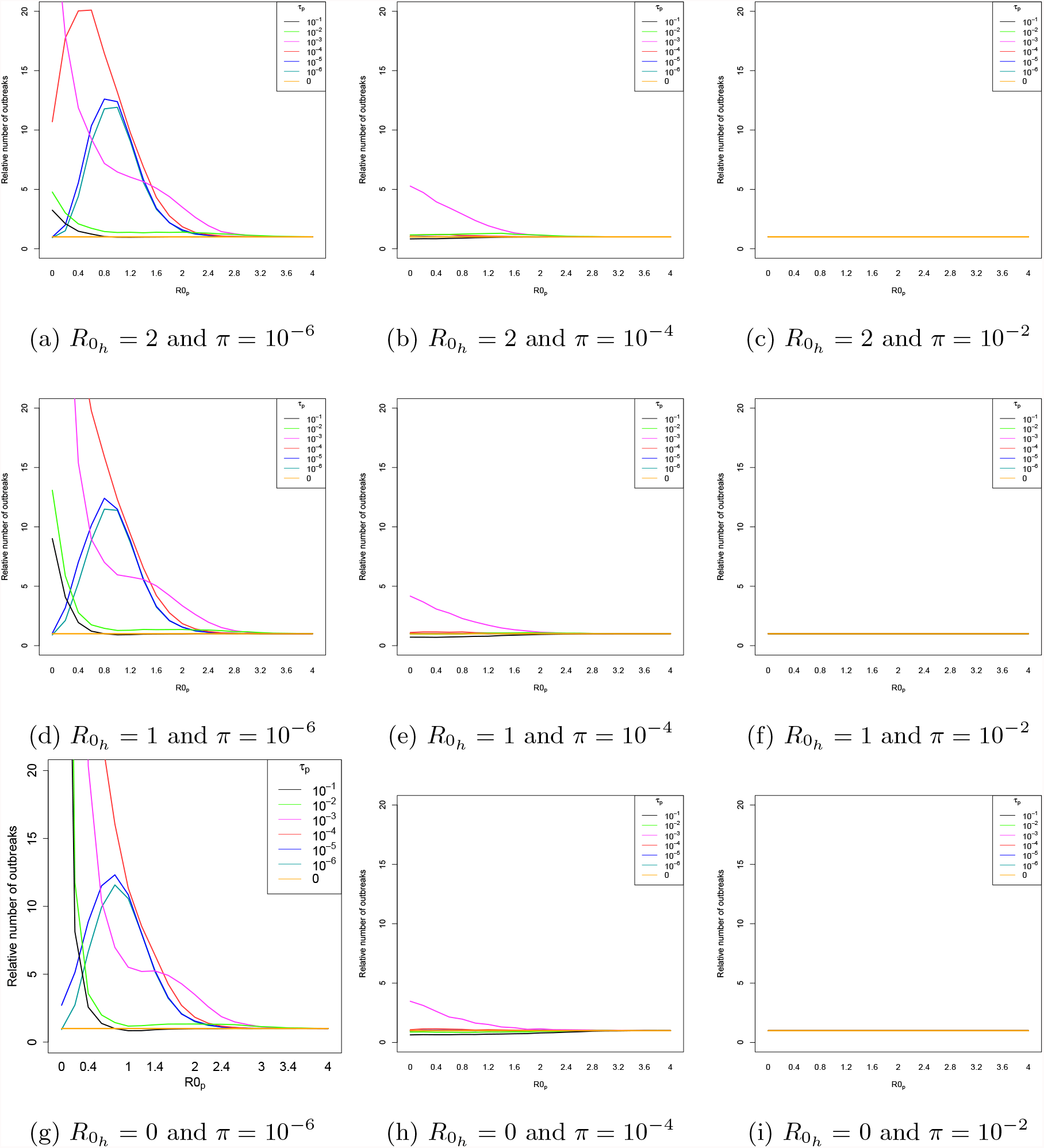
Relative number of outbreaks between the full model and the reservoir model. A low, intermediate and high value of the transmission between individuals in the intermediate host (respectively 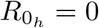, 1 and 2) and inter-incidental transmission (respectively *π* = 10^−6^, 10^−4^ and 10^−2^) are considered. The relative number of outbreaks is depicted as a function of the transmission between individuals in the target population 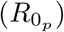. Each curve represents a spillover transmission from the reservoir to the target population (*τ*_*p*_) ranging from 10^−6^ to 10^−1^. A high spillover rate from the reservoir to the intermediate host is considered *τ*_*h*_ = 10^−2^.

#### D.2 The relative size of the largest outbreak

**Figure D.4:**
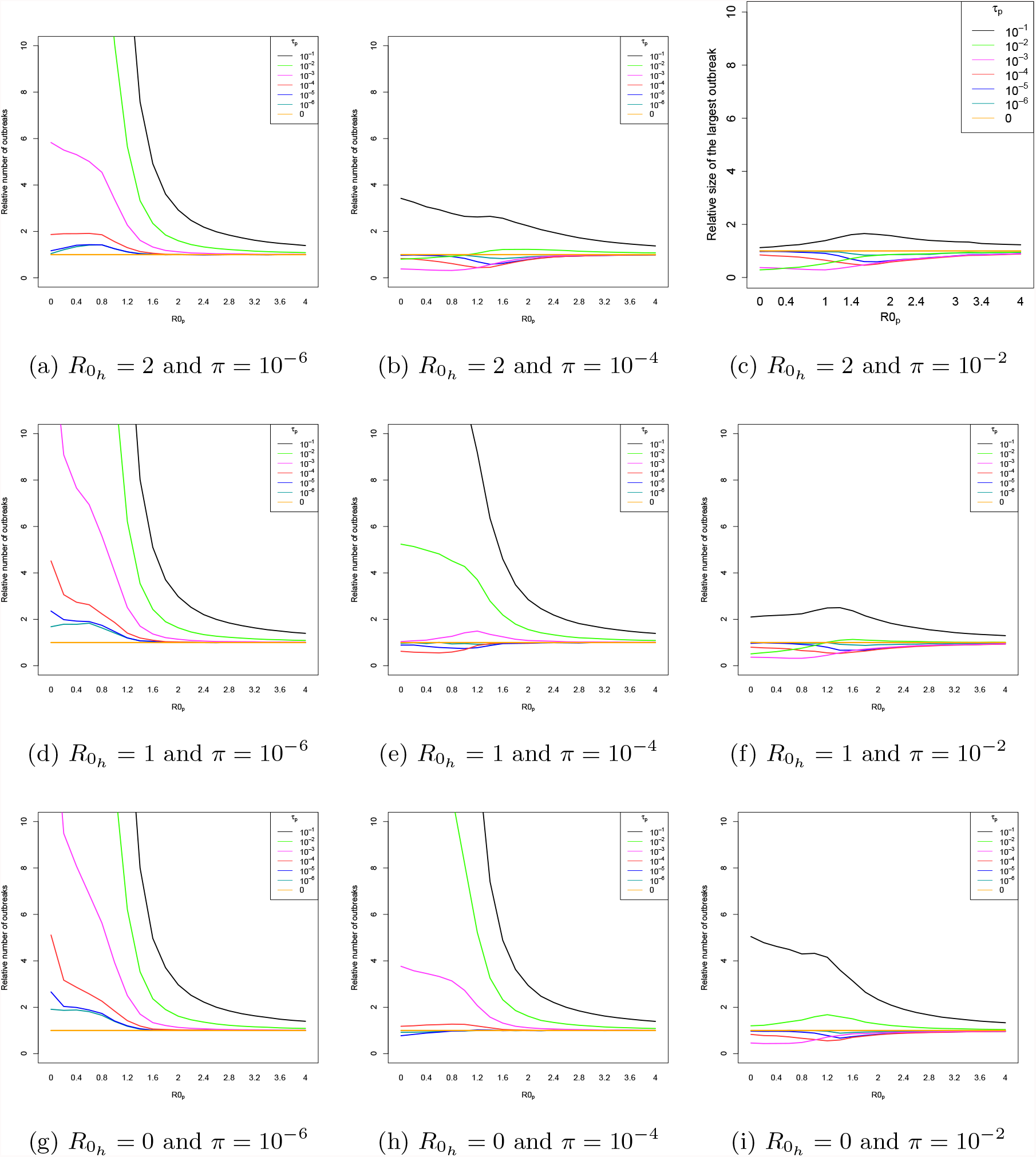
Relative size of the largest outbreak between the full model and the reservoir model. A low, intermediate and high value of the transmission between individuals in the intermediate host (respectively 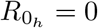, 1 and 2) and inter-incidental transmission (respectively *π* = 10^−6^, 10^−4^ and 10^−2^) are considered. The relative size of the largest outbreak is depicted as a function of the transmission between individuals in the target population 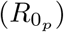. Each curve represents a spillover transmission from the reservoir to the target population (*τ*_*p*_) ranging from 10^−6^ to 10^−1^. A low spillover rate from the reservoir to the intermediate host is considered *τ*_*h*_ = 10^−6^.

**Figure D.5:**
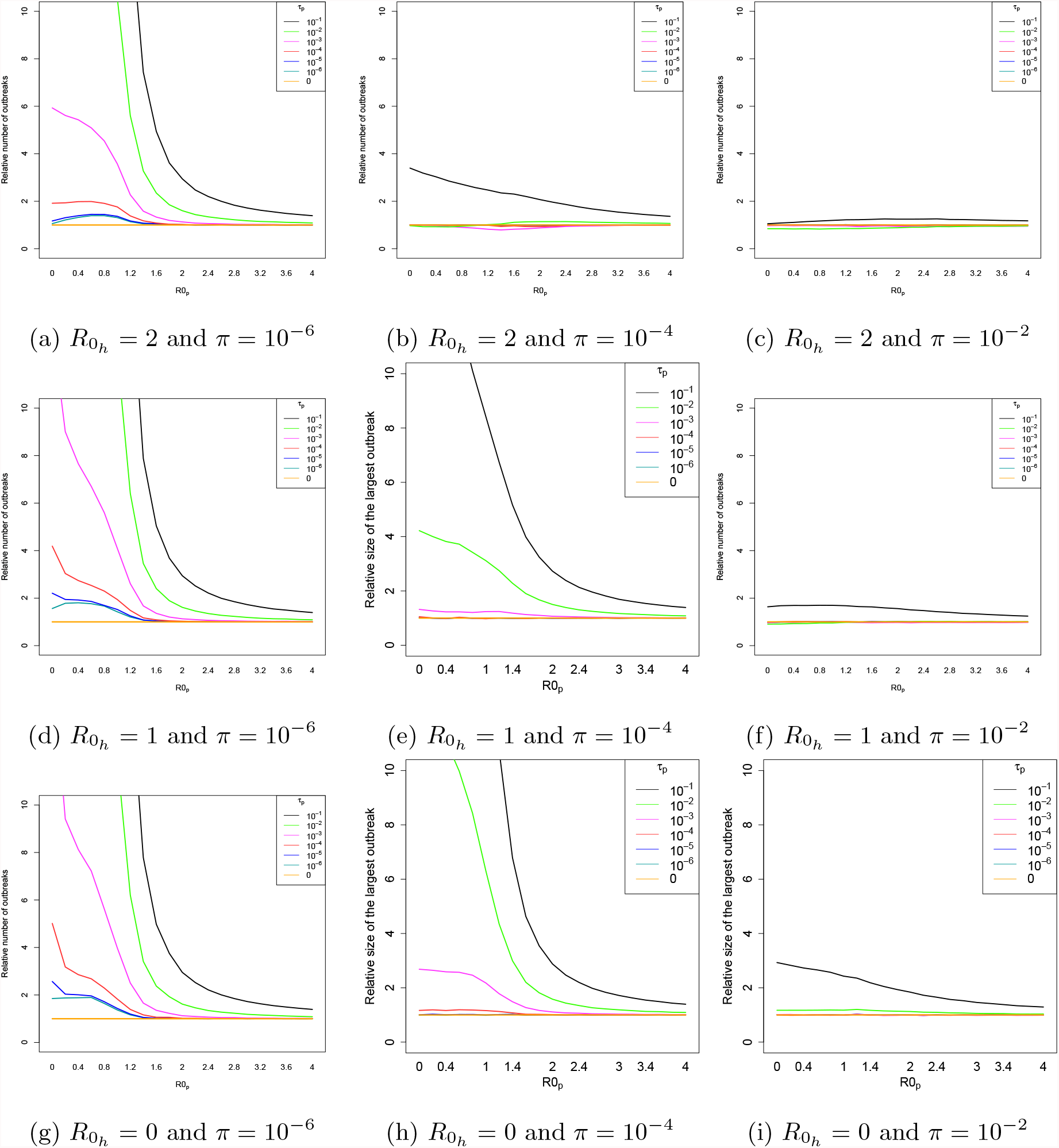
Relative size of the largest outbreak between the full model and the reservoir model. A low, intermediate and high value of the transmission between individuals in the intermediate host (respectively 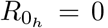, 1 and 2) and inter-incidental transmission (respectively *π* = 10^−6^, 10^−4^ and 10^−2^) are considered. The relative size of the largest outbreak is depicted as a function of the transmission between individuals in the target population 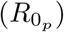. Each curve represents a spillover transmission from the reservoir to the target population (*τ*_*p*_) ranging from 10^−6^ to 10^−1^. An intermediate spillover rate from the reservoir to the intermediate host is considered *τ*_*h*_ = 10^−4^.

**Figure D.6:**
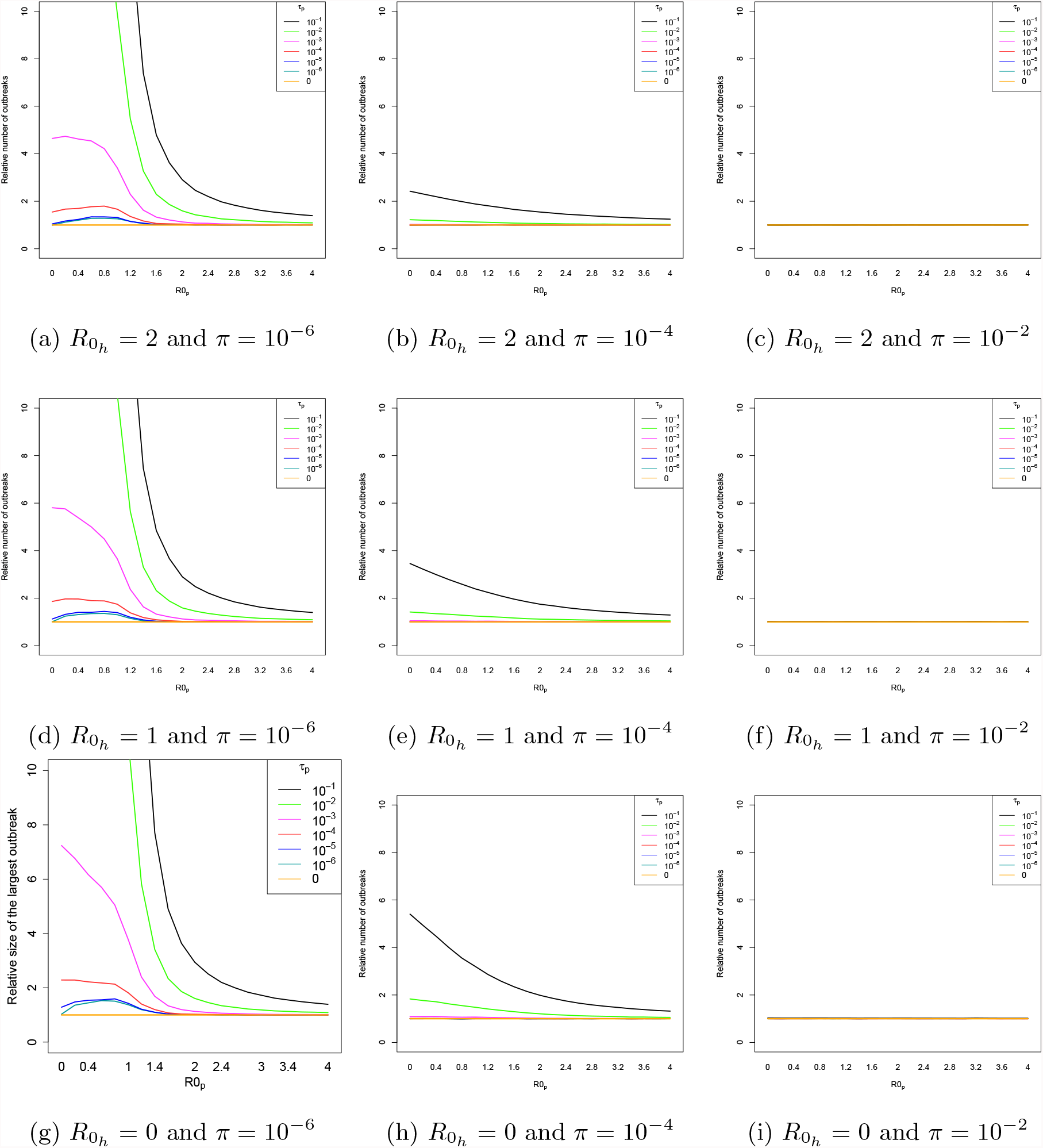
Relative size of the largest outbreak between the full model and the reservoir model. A low, intermediate and high value of the transmission between individuals in the intermediate host (respectively 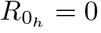, 1 and 2) and inter-incidental transmission (respectively *π* = 10^−6^, 10^−4^ and 10^−2^) are considered. The relative size of the largest outbreak is depicted as a function of the transmission between individuals in the target population 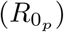. Each curve represents a spillover transmission from the reservoir to the target population (*τ*_*p*_) ranging from 10^−6^ to 10^−1^. A high spillover rate from the reservoir to the intermediate host is considered *τ*_*h*_ = 10^−2^.

